# Integrative modeling of tumor DNA methylation identifies a role for metabolism

**DOI:** 10.1101/057638

**Authors:** Mahya Mehrmohamadi, Lucas K. Mentch, Andrew G. Clark, Jason W. Locasale

## Abstract

DNA methylation varies across genomic regions, tissues and individuals in a population. Altered DNA methylation is common in cancer and often considered an early event in tumorigenesis. However, the sources of heterogeneity of DNA methylation among tumors remain poorly defined. Here, we capitalize on the availability of multi-platform data on thousands of molecularly-and clinically-annotated human tumors to build integrative models that identify the determinants of DNA methylation. We quantify the relative contribution of clinical and molecular factors in explaining within-cancer (inter-individual) variability in DNA methylation. We show that the levels of a set of metabolic genes involved in the methionine cycle that are constituents of one-carbon metabolism are predictive of several features of DNA methylation status in tumors including the methylation of genes that are known to drive oncogenesis. Finally, we demonstrate that patients whose DNA methylation status can be predicted from the genes in one-carbon metabolism exhibited improved survival over cases where this regulation is disrupted. To our knowledge, this study is the first comprehensive analysis of the determinants of methylation and demonstrates the surprisingly large contribution of metabolism in explaining epigenetic variation among individual tumors of the same cancer type. Together, our results illustrate links between tumor metabolism and epigenetics and outline future clinical implications.

## Introduction

DNA methylation is a major epigenetic mechanism that determines cellular outcome by regulating gene expression and chromatin organization^1^ in a fashion more dynamic than previously appreciated^2^. Altered DNA methylation is frequently observed in cancers compared to corresponding normal cells^3–7^. For example, global DNA hypomethylation^8^ and tumor suppressor silencing by DNA hypermethylation are two of the most well characterized cancer associated alterations common across many human malignancies^9–11^. In addition to hypo and hyper methylation, cancer cells exhibit increased variability in DNA methylation across large portions of the genome compared to their corresponding normal tissues^12^. A previous study showed that, for several cancer types, variation in methylation levels among tumor samples is significantly higher than normal samples of the same tissue of origin^4^, possibly indicating that deregulated epigenetics provides tumor cells with potential proliferative advantages^6^. While inter-tissue variability in DNA methylation is mainly explained by differentiation and tissue-specific regulatory mechanisms^13, 14^, very little is known about the functions and determinants of the high inter-individual variation among tumors of the same tissue type. Notably, a recent twin study on the determinants of inter-individual variability in DNA methylation reported that genetic difference among individuals account for only 20% of total variance with the remaining variance explained by environmental and stochastic factors that are yet to be identified^15^.

The source of the methyl group for methylation is S-adenosylmethionine (SAM) which is generated from the methionine (met) cycle and is coupled to serine, glycine, one-carbon (SGOC) metabolism^16^. A large body of evidence indicates numerous roles for one-carbon metabolism in proliferation and survival of tumor cells through its roles in biosynthesis and redox metabolism^16–19^. The met cycle also mediates histone and DNA methylation in physiological conditions and provides a link between intermediary metabolism and epigenetics^20–22^. Although the network contributes methyl units to DNA, whether and to what extent this interaction is apparent in tumors and may contribute to cancer biology is unknown.

We set out to comprehensively quantify the contribution of various factors in explaining variation in DNA methylation. The advent of standardized genomics and other high-dimensional multi-platform ‘omics’ data through The Cancer Genome Atlas (TCGA) allows for systematic assessments of molecular features across cancers^23^. With combined statistical analysis, computational modeling, and machine-learning approaches, we directly evaluated the quantitative contributions of molecular and clinical variables that lead to DNA methylation. We found a surprisingly large contribution for the expression of the methionine cycle and related SGOC network genes in determining DNA methylation and identified numerous contexts where this interaction may contribute to cancer pathology.

## Results

### Integrative modeling allows for quantitation of determinants of variation in DNA methylation

It has been previously proposed that factors normally regulating the epigenome are disrupted in cancer, leading to increased variability of the cancer epigenome^6^. However, the nature and contributions of such factors is largely unknown. Upon analysis of global and local DNA methylation in tumors as measured by the Illumina Infinium HumanMethylation450K BeadChip arrays, we indeed found higher variation among tumors from the same tissue vs. between different tissue types (Supplementary Information; Supplementary Fig. 1a-d; Online Methods). Arrays were used over bisulfite sequencing because of the higher availability of these data in a standardized format allowing for an integrative analysis. To establish quantitative relationships between DNA methylation and molecular and clinical features of tumors, we developed an integrative statistical modeling and machine-learning approach with the goal of identifying the relative contributions to within-cancer DNA methylation variation (Online Methods). We incorporated hundreds of variables into comprehensive statistical models of DNA methylation (Fig. 1a). Factors with a known role in DNA methylation machinery (chromatin remodeling enzymes and transcription factors), as well as factors with a potential biochemical link to DNA methylation (SAM-metabolizing enzymes, met cycle enzymes, and other serine, glycine, one-carbon (other SGOC) enzymes that are connected to the met cycle^24^) were together considered (Fig. 1a). We also curated available clinical information such as age, gender, and cancer stage in the calculations where appropriate. Furthermore, since mutations are known to affect the cancer methylome^25^, we included all recurrent genetic lesions (somatic mutations and copy number alterations) for each cancer type in our models. Together, over 200 variables were collectively analyzed for each cancer type (see Supplementary Table 1). Our models are therefore not completely agnostic as we pre-select classes of biological variables that are known to affect DNA methylation to avoid loss of statistical power by including too many features (e.g. expression of all genes in the genome). Therefore, to test for potential bias, we also considered the expression levels of sets of random genes with functions non-related to DNA methylation as additional variables in our models (see Online Methods).

Subsequently, we incorporated all variables into unbiased selection algorithms suitable for dealing with large numbers of prediction variables. For this task, we considered two independent approaches: a generalized linear model (Elastic Net)^26, 27^ and a machine-learning algorithm (Random Forest)^28, 29^. A distinct computation was carried out for each 10kilobase (kb) genomic region with variable methylation (sd>0.2) in each cancer type. Samples of each cancer type were divided into three independent test subsets and three training subsets and separate models were generated using each subset. The models were then combined resulting in a single final model for each 10kb region of DNA methylation in each cancer. Model performance was evaluated by measuring mean squared prediction error of test samples from Elastic Net and Random Forest separately (Online Methods).

**Figure 1.**
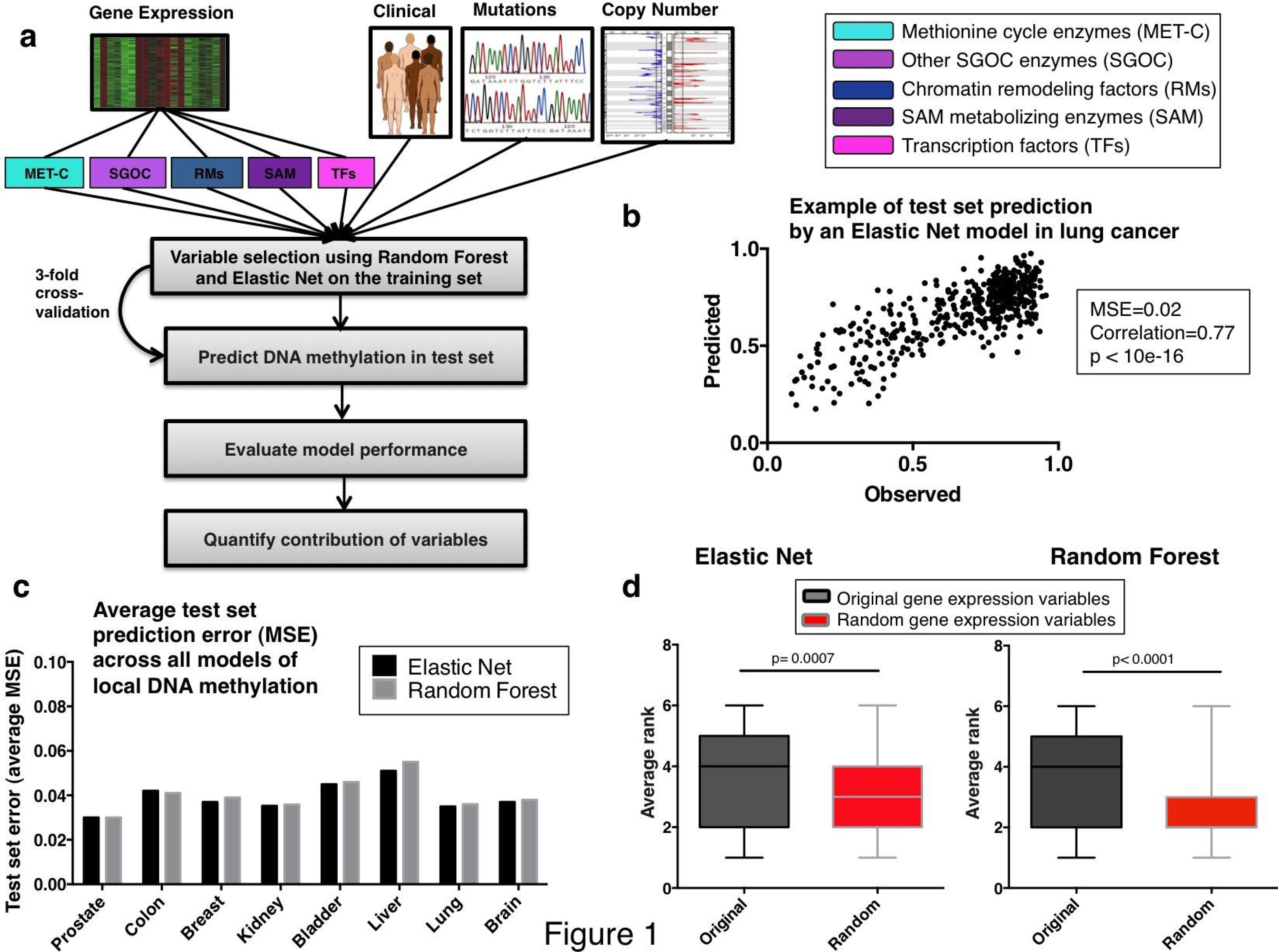
Integrative modeling of local DNA methylation levels.

a) Schematic summarizing the integrative approach utilized for modeling local DNA methylations. DNA methylation at a given 10kb region was predicted by incorporating relevant gene expression, somatic mutation, copy number alteration, and clinical information into integrative models (see Supplementary Table 1 for the complete list of variables included for each cancer type).

b) An example of an Elastic Net model performance in lung cancer. The x-axis shows true values of DNA methylation in each sample, and the y-axis shows the value predicted by the integrative modeling in the same sample when it was in the test subset.

c) Summary of overall model performance. For each cancer, the mean squared errors of test set predictions by Elastic Net and Random Forest were averaged across all models of local DNA methylation.

d) Comparison of original gene expression variables with randomly selected variance-matched genes. The y-axis shows the average rank of each gene expression category based on average variable usage score across all Elastic Net models (left) and average variable importance score across all Random Forest models (right) of local DNA methylation in brain cancer. P-values associated with the Mann-Whitney test between the ranks across all models are shown (see Online Methods).

We observed that our models predicted test set DNA methylation with small mean squared error (MSE < 0.04) in many regions across the genome (Fig. 1b). Comparison of the performances of the two methods showed that Random Forest and Elastic Net algorithms were able to predict DNA methylation with comparable MSEs on average (Fig. 1c; Supplementary Fig. 2a). In general, predictability of local DNA methylation was largely dependent on cancer type as well as chromatin region in each model. For example, we observed that local DNA methylation was most predictable in prostate and lung cancers and least predictable in liver and bladder cancers (Fig. 1c; Supplementary Fig. 2a). Together with the high variation in local DNA methylation levels seen in liver and bladder cancers (Supplementary Fig. 1d), these results suggest a higher stochasticity in the epigenetic signatures for these two cancer types compared to others in this study.

Upon annotating genomic regions where local DNA methylation could be predicted with a low error (MSE < 04) in each cancer, we found that the majority of the predictable regions lie within 20kb of the transcriptional start site (TSS) of a gene (Supplementary Fig. 2b), suggesting that regulation of DNA methylation by the factors included in our models is stronger at genic regions.

We next performed a set of tests to evaluate the robustness of our modeling approach. To this end, we compared the original gene expression variables included in our models, with a group of variance-matched randomly selected genes from the genome (see Online Methods) in their ability to predict DNA methylation. In the presence of both groups of gene expression variables (original and random), both Elastic Net and Random Forest models selected our original variables significantly more frequently than random genes (Mann-Whitney p-value = 0.0007 for Elastic Net and < 0.0001 for Random Forest) (Fig. 1d; see Online Methods). These results validate our models and confirm that the Elastic Net and Random Forest algorithms are suitable for quantitation of variable contributions in determining DNA methylation.

### Metabolism is a major predictor of DNA methylation in human cancers

Using the results of the integrative modeling, we next quantified the relative contribution of different functional classes of variables in explaining DNA methylation variation within each cancer type. For this, we measured two independent metrics, one using the Random Forest variable importance scores, and the other using a binary score for whether or not a variable was selected by the Elastic Net models (non-zero co-efficient). For each variable, an overall importance score was calculated by averaging its relative importance across all models of 10kb DNA methylations, and an overall usage score was calculated by measuring the fraction of 10kb regions in which Elastic Net models selected the variable (Online Methods). To estimate the contribution of each functional class of variables in explaining total variation in DNA methylation, we pooled all variables in the same functional category and averaged across their importance and usage scores separately (Supplementary Fig. 2c,d).

Results from both Random Forest and Elastic Net algorithms identified a considerable contribution from the variables within the SGOC metabolic network relative to other classes of variables (“Other SGOC enzymes” was the 2^nd^ highest scoring among all classes, closely following “Transcription factors” according to both methods. “Methionine cycle enzymes” was the 3^rd^ and 4^th^ according to Random Forest and Elastic Net, respectively) (Fig. 2a). Previous studies have shown that transcription factor abundance and occupancy strongly mediate dynamic DNA methylation turnover in regulatory regions^30, 31^. Consistent with this observation, our results confirm the “Transcription factors” class has the highest contribution to predicting DNA methylation levels across human tumors. Notably, even in the presence of most if not all known variables that are thought to mediate the status of DNA methylation, metabolic factors still uniquely explained a large part of the variability in methylation (Fig. 2a).

**Figure 2.**
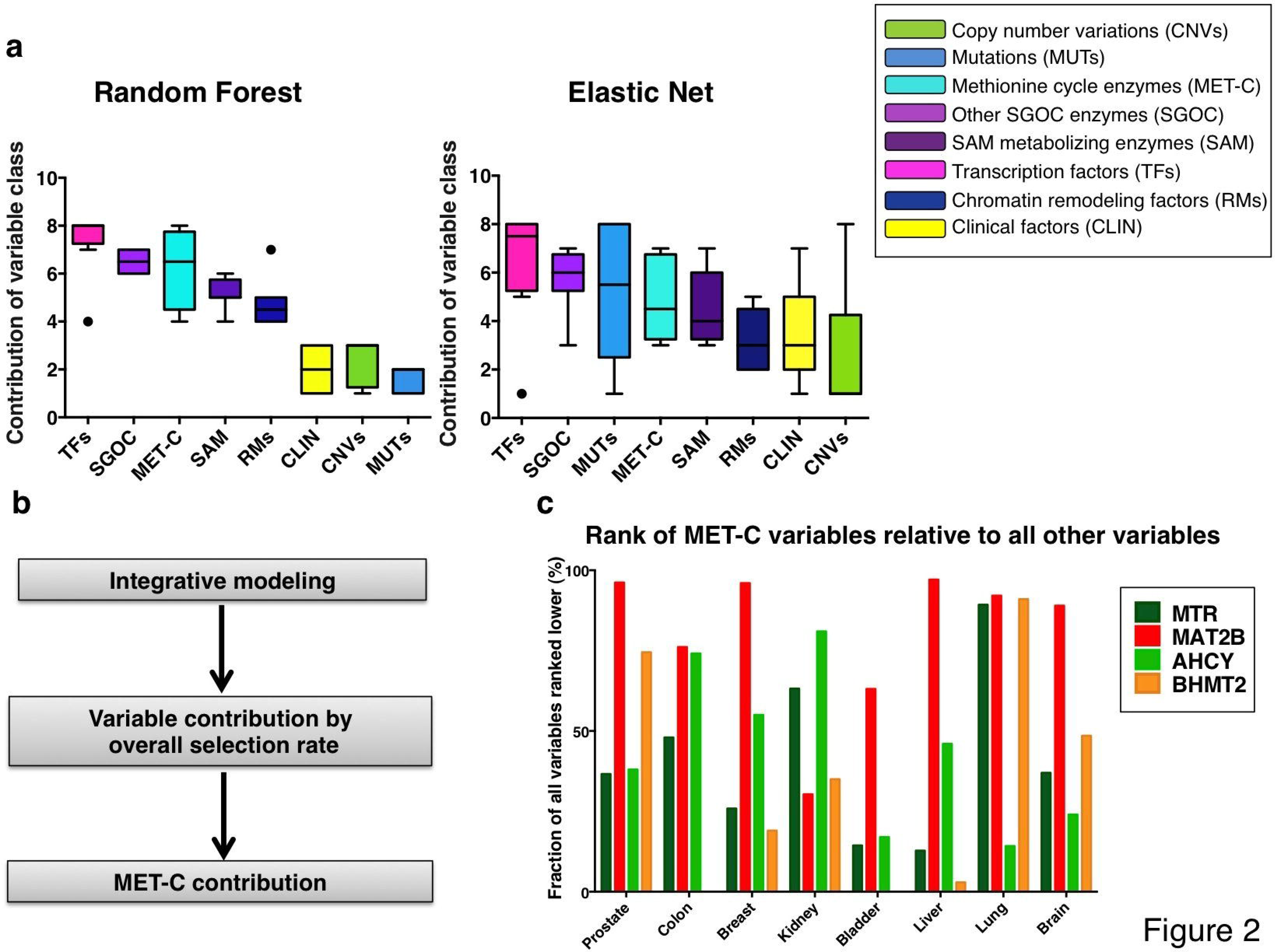
Contribution of different functional classes of variables to DNA methylation variation.

a) Relative contributions of the variable classes according to Random Forest average variable importance (left), and Elastic Net average variable usage (right) are shown averaged across all cancers (Online Methods). The y-axis shows the average rank of each class across cancers (with higher values corresponding to higher contribution).

b) Diagram summarizing the steps taken toward calculating overall contribution of each of the met cycle variables relative to other variables in explaining variability local DNA methylations.

c) Ranking all variables according to their overall selection rate (usage) across all models of local DNA methylation in each cancer. The y-axis shows the percent of variables that ranked lower than each of the met cycle variables (i.e. made less contribution to DNA methylation) in each cancer (BHMT2 was removed from the models of colon and bladder cancers due to low expression).

Given the contribution of the methionine cycle and its biochemical link to DNA methylation, we further explored the variables within the met cycle class compared to all other variables in their ability to predict DNA methylation (Fig. 2b). Within the met cycle class, methionine adenosyltransferase 2 beta (MAT2B) and betaine-homocysteine S-methyltransferase 2 (BHMT2) exhibited higher predictive values than methionine synthase (MTR) and adenosylhomocysteinase (AHCY) on average (Supplementary Fig. 2e,f). Notably, in the presence of the nearly 200 other variables in the computations, the met cycle — especially MAT2B — still contributed substantially to DNA methylation prediction (MAT2B was ranked among the top 5% of highly selected variables in prostate, breast, liver, lung, and brain cancers) (Fig. 2c). We observed that the levels of MAT2B contribute to DNA methylation in nearly half of the variable regions across the genome even after accounting for various factors related to DNA methylation (MAT2B was selected by 42% of all Elastic Net models with MSE < 0.04 on average) (Supplementary Fig. 2e). Together, our results confirm that metabolism contributes to DNA methylation in many cases of human cancer and the association between metabolism and DNA methylation is stronger in some genomic regions than others.

### Functional annotation of metabolically regulated regions of DNA methylation

Results of the integrative modeling across cancers indicate that defined regulation of DNA methylation happens in regions where gene expression may be affected, thereby suggesting that this regulation could drive essential cancer biology. We next set out to characterize all regions across the genome where the association between DNA methylation and the met cycle activity is particularly strong. To identify such regions, we designed a scanning algorithm to locate genomic regions spanning multiple CpGs with significant peaks of correlation of methylation with expression of met cycle enzymes (Fig. 3a; Online Methods). We performed this analysis on each of the eight cancer types separately and identified distinct peak sets across the genome (Supplementary Table 2). To assess potential bias toward highly methylated regions and regions where there is higher probe density, we analyzed the relationship between average absolute methylation of individual CpGs and their correlation with met cycle expression, and found no significant association (p-value of correlation = 0.62), confirming that the identified peaks are distinct from highly methylated regions (Fig. 3b; Online methods).

**Figure 3.**
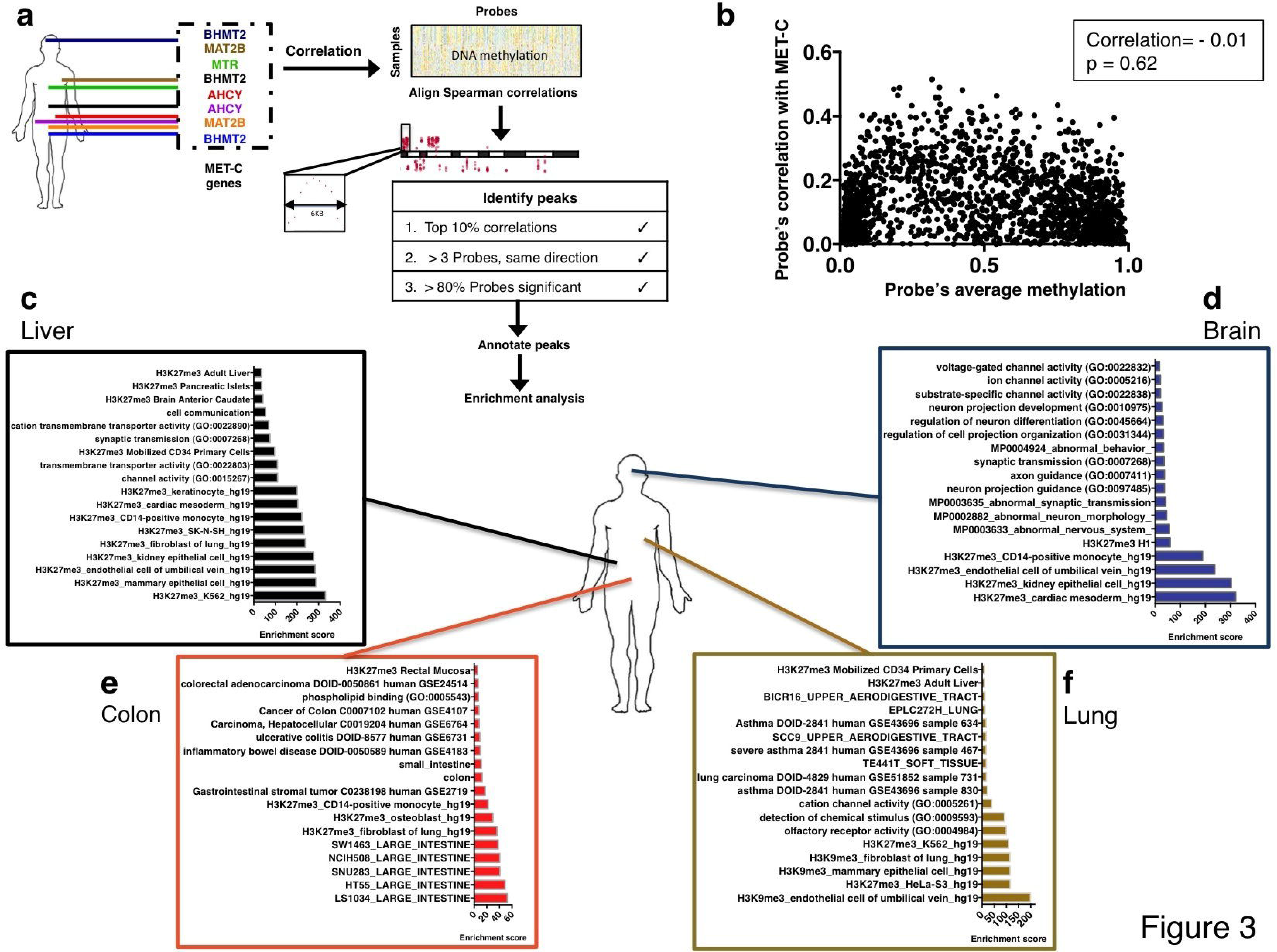
Genome-wide screening for metabolically regulated regions.

a) Schematic describing the algorithm used for finding genomic regions where DNA methylation might be regulated by the met cycle (see Online Methods).

b) Assessment of the relationship between met cycle correlation and absolute methylation. The y-axis shows the Spearman rho for correlation of 2000 randomly selected probes with the expression of AHCY in colon cancer. The x-axis shows the average methylation level of the same probes across the colon cancer samples in the study (see Online Methods).

c-f) Pathway enrichment analyses of genes overlapping peaks. Results are depicted by functional annotation of genes located within peaks of correlation between met cycle and DNA methylation (see Online Methods for description of gene sets and enrichment scores; see Supplementary Fig. 6 for additional cancer types).

Density plots of peak distributions relative to the TSS of the nearest gene were concentrated around the TSS in all cancers (Supplementary Fig. 3a), as expected given the higher density of probes in gene regulatory regions in the Illumina arrays (Supplementary Fig. 3b). However, by further visualizing the distribution of the peaks immediately surrounding the TSS, we observed that peak distributions are more diffuse around the TSS (Supplementary Fig. 3c) compared to the probe density distribution control (Supplementary Fig. 3d). This suggests potential enrichment in areas of the genome overlapping with gene body regions and CpG island shores where dysregulated DNA methylation has previously been observed in human cancers^6^. The peak distribution density plots extended up to a few hundred kilobases in distance from the nearest TSS, suggesting that DNA methylation at inter-genic parts of the genome may also be affected by the activity of met cycle.

We next tested the met cycle specificity of the identified peaks by correlating them with expression of randomly selected genes in the genome (Online Methods; Supplementary Fig. 4a). For the majority (>83%) of the identified peaks, the met cycle's correlation with DNA methylation was significantly non-random (p-value < 0.05) (Supplementary Fig. 4b). These results show that our approach was able to identify genomic regions where DNA methylation levels are specifically affected by the met cycle activity.

We next set out to identify genes that overlap with the identified peaks in each cancer type. Functional annotation of genes overlapping these peaks by means of pathway enrichment analyses across a comprehensive collection of more than 70 gene-set libraries^32^ showed enrichment of epigenetic features in these regions consistently across all cancers. Strikingly, many of our peaks overlapped with peaks of histone-3 lysine-27 tri-methylation (H3K27me3) (Fig. 3c–f; Supplementary Fig. 5a-d) as reported by both the encyclopedia of DNA elements (ENCODE) human project^33^ and the RoadMap epigenomics project^34^. In cancers of the lung and bladder, histone-3 lysine-9 tri-methylation (H3K9me3) peaks were also significantly enriched (Fig. 3f; Supplementary Fig. 5c). H3K27me3 and H3K9me3 are both associated with repression of gene expression^35^. Our findings therefore suggest that variation in the met cycle's activity may contribute to aberrant expression from normally silenced loci and heterochromatin instability in cancer.

In addition to histone marks, tissue-specific and cell identity gene sets were also enriched in relevant cancer types, including “breast and ovarian cancer genes” in breast cancer (Supplementary Fig. 5a); “abnormal nervous system” and “abnormal neuron morphology” in brain cancer (Fig. 3d); “asthma” and “lung carcinoma” gene sets in lung cancer (Fig. 3f); “kidney-specific” gene set in kidney cancer (Supplementary Fig. 5b); and “large intestinal genes”, “inflammatory bowl disease”, and “colorectal carcinoma” gene sets in colon cancer (Fig. 3e). Finally, a number of developmental and signaling pathways were among the enriched pathways including “TGF-beta signaling” in kidney (Supplementary Fig. 5b), “cell communication” pathway in liver (Fig. 3c), and “G-protein coupled signaling” in bladder cancer (Supplementary Fig. 5c). Organ and embryonic morphogenesis pathways were enriched in breast (Supplementary Fig. 5a), bladder (Supplementary Fig. 5c), and prostate (Supplementary Fig. 5d), all of which are hormonally driven cancers. Interestingly, a previous study in breast cancer showed that embryonic developmental genes are enriched in regions of DNA hypomethylation compared to normal breast^36^. Together, these results illustrate the functional importance of the relationship between met cycle and DNA methylation across cancers.

### Contribution of metabolism to DNA methylation at cancer gene loci

So far, we have shown that there is a surprising enrichment of peak regions of metabolically regulated DNA methylation at loci that link to essential aspects of cell identity and chromatin structure. We next questioned whether cancer-specific loci may also exhibit this interaction. We chose 19 well-characterized cancer-related genes such as *TP53, PTEN*, and *ESR1*, as well as 4 genes frequently differentially methylated in cancer *APC, RASSF1, GSTP1*, and *MGMT* (Online Methods). A recent study showed that DNA methylation for any given gene has two major principal components: one representing the promoter region and the other representing the coding sequence^37^. Furthermore, CpG methylation at promoter regions of genes is typically associated with repression, while gene body methylation is thought to increase expression^38^. We therefore applied our integrative modeling to DNA methylation at promoter and gene body regions of each cancer gene separately. In addition to the integrative approach, we also generated models using only the met cycle genes as prediction variables to quantify the predictive ability of met cycle in the absence of other factors. Thus, each cancer gene locus was analyzed once using the integrative approach and once using met cycle alone and 3-fold cross validation was performed in each case as previously described (Online Methods). Model performance was evaluated by calculating the error of prediction of test set methylation, as shown for two examples in Fig. 4: estrogen receptor *(ESR1)* promoter in breast cancer (MSE = 0.004) (Fig. 4a), and androgen receptor (AR) promoter in prostate cancer (MSE = 0. 001) (Fig. 4b). *ESR1* promoter methylation in breast cancer and *AR* promoter methylation in prostate cancer are two examples of events that are known to contribute to the pathogenesis and prognosis of the corresponding tumor types ^39–42^. We further assessed the integrative models of promoter methylation at these two loci, and found many SGOC (including met cycle) variables among the top predictive variables of promoter methylation according to the variable importance measures (Fig. 4c,d; Online Methods).

**Figure 4.**
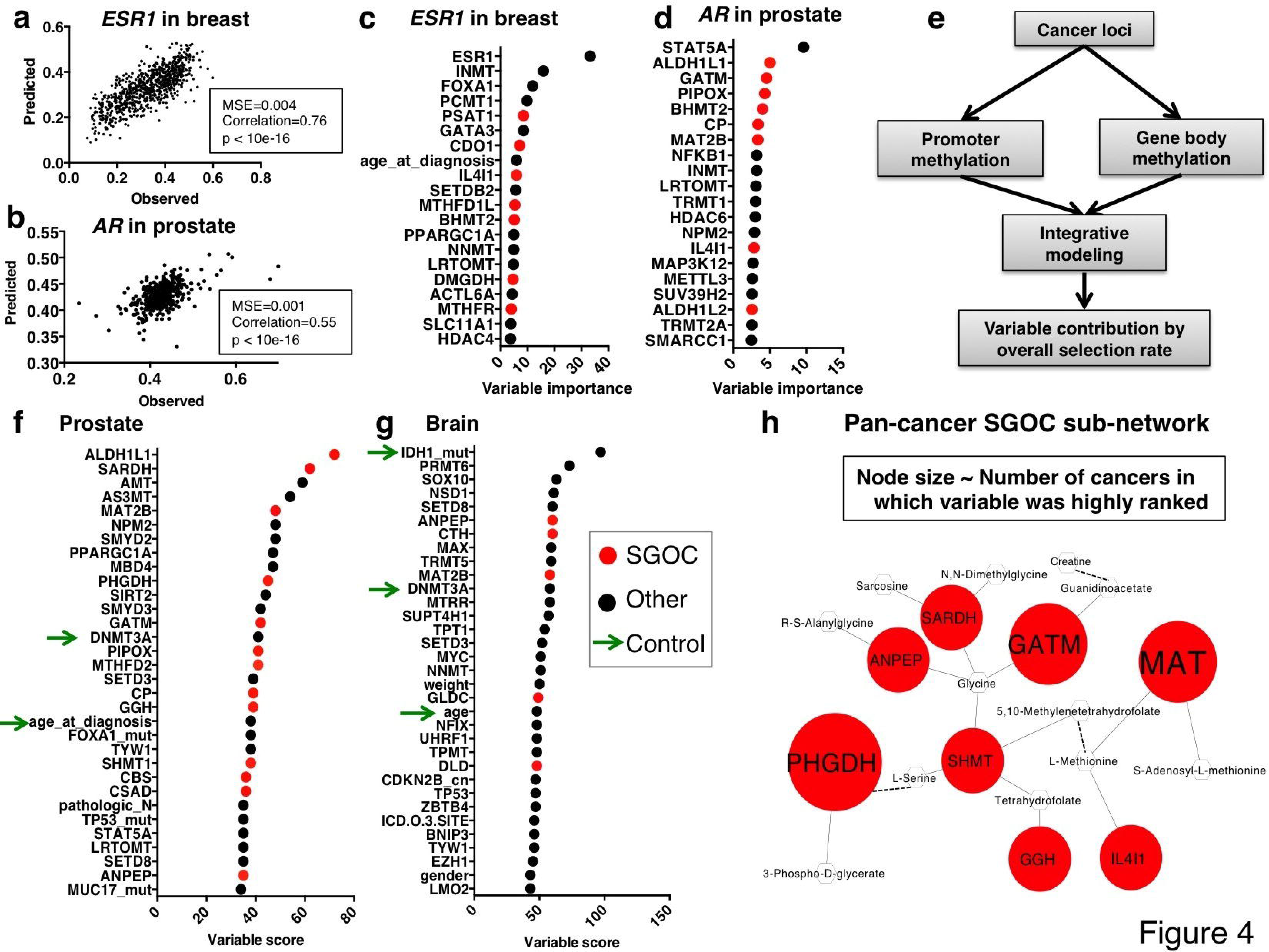
Contribution of metabolism to DNA methylation at cancer loci.

a) Prediction of ESR1 promoter methylation in test set samples in breast cancer. The x-axis shows the true methylation value at ESR1 promoter for the test set samples, while the y-axis shows the corresponding values as predicted by Elastic Net.

b) Prediction of AR promoter methylation in test set samples in prostate cancer. The x-axis shows the true methylation value at AR promoter for the test set samples, while the y-axis shows the corresponding values as predicted by Elastic Net.

c-d) Top 20 variables as ranked based on the variable importance score from Random Forest model of ESR1 promoter methylation in breast cancer (c), and AR promoter methylation in prostate cancer (d). Variables in the serine, glycine, one-carbon (SGOC) network (including the met cycle enzymes and other SGOC enzymes) are shown in red and all other variables are shown in black.

e) Schematic depicting the ranking of all variables based on the combined results from promoter and gene body methylation at the 23 cancer loci in the study.

f-g) Variables that were most predictive of cancer gene methylation on average (top 15%) are listed and ranked in order of increasing contribution (variable score = percent variable usage by Elastic Net averaged across all models of cancer gene body and promoter methylation). Variables in the serine, glycine, one-carbon (SGOC) network (including the met cycle genes and other SGOC genes) are shown in red and all other variables are shown in black. Green arrows point to previously published factors associated with variations in DNA methylation in each cancer type (positive controls). (Variable names: official gene symbols are used to show gene expression variables (including “Methionine cycle enzymes”, “Other SGOC enzymes”, “Transcription factors”, “Chromatin remodeling factors”, and “ SAM metabolizing enzymes”), while “_mut” and “_cn” suffixes following gene symbols denote “Mutations” and “Copy number variations”, respectively. For “Clinical factors”, variable names match the descriptors used in the TCGA clinical data files) (see Supplementary Fig. 9 for additional cancer types).

h) Sub-network of SGOC genes contributing to DNA methylation in multiple cancer types (at least 4 cancers based on Elastic Net models and at least 3 cancers based on Random Forest models). Red nodes represent genes while smaller white nodes represent metabolites. Solid edges denote direct biochemical links and dashed edges denote indirect biochemical links through enzymatic reactions not shown. Node sizes for the gene nodes correspond to the number of cancer types wherein each enzyme contributed significantly to cancer gene methylation. (Node size scores: phosphoglycerate dehydrogenase (PHGDH) = 6, MAT (including MAT2B and MAT2A) = 5, glycine amidinotransferase (GATM) = 5, serine hydroxymethyltransferase 1 and 2 (SHMT) = 4, sarcosine dehydrogenase (SARDH) = 4, alanyl aminopeptidase (ANPEP) = 4, L-amino acid oxidase (IL4I1) = 4, gamma-glutamyl hydrolase (GGH) = 4).

Notably, the models across all cancers in the study were able to predict cancer gene methylation with high accuracy even using the met cycle variables in the absence of all other variables (85% of the predictions were made with MSE < 0.01) (Supplementary Fig. 6a,b). As in the case of local methylation, cancer gene methylation was also more strongly explained by the expression of MAT2B compared with other met cycle variables on average (selected by 24% of all integrative models) (Supplementary Fig. 6c), consistent with the function of this enzyme that directly affects SAM levels. Relative variable class comparisons confirmed considerable contribution from the “Methionine cycle enzymes” and “Other SGOC enzymes” among other classes of variables (highest after “Transcription factors” and “Mutations”) (Supplementary Fig. 6d,e).

We independently evaluated these findings by applying the same models to both permuted cancer gene methylation values and also randomly generated methylation values (Supplementary Fig. 7a). In all tests, met cycle contribution was significantly (p-value < 10e-16) higher when applied to cancer gene methylation vs. permutations or random numbers (Online Methods; Supplementary Fig. 7b), confirming the specificity of signals contained in the true DNA methylation values at cancer loci. Furthermore, we tested the performance of the machine-learning algorithm using randomly generated variables for prediction of cancer gene methylation (Online Methods) and found in each of the cases tested, that the predictions made with the original variables are uniformly more accurate than what is made using simulated random variables (original model MSE smaller by 1.4-2 fold than random model MSE on average) (Supplementary Fig. 8a-d). We also simulated a dataset where prediction variables and the response are related via linear relationships and compared the accuracy of predictions in this simulated linear dataset with our original dataset (Online Methods). We saw in all cases that the improvement in MSE from our dataset (MSE 1.4-2 fold smaller than random MSE) is even more than what we observed with data of the same dimension that have a linear relationship (MSE 1.3 fold smaller than random MSE) (Supplementary Fig. 8e,f). These independent tests confirm that machine-learning using the Random Forest algorithm is able to identity non-random signals in the data, and also that it can detect non-linear relationships between prediction variables and the response.

Next, we ranked all of the variables based on their overall usage according to the integrative models of cancer gene promoter and body methylations (Fig. 4e). Notably, many SGOC (including met cycle) enzymes were among the most frequently selected variables in all cancers (Fig. 4f,g; Supplementary Fig. 9a-f). Importantly, our models highly ranked many clinical and molecular factors previously shown to be associated with DNA methylation in the existing literature (green arrows in Fig. 4f, g and Supplementary Fig. 9a-f). Examples of such positive controls include DNA methyltransferase (DNMT3A or DNMT3B) enzymes^43^ that were consistently among the top variables in all cancers (Fig. 4f,g; Supplementary Fig. 9a-f), and patient's age (or age at diagnosis)^15, 44^ that was highly ranked in prostate, colon, breast, kidney, and brain (Fig. 4f,g; Supplementary Fig. 9a-f). We also observed ER-status to be one of the most important contributors to DNA methylation variation in breast cancer consistent with previous publications^45^ (Supplementary Fig. 9b). Furthermore, we found the mutational status of the histone methyltransferase SET-domain containing-2 (*SETD-2*) as a significant contributor in kidney (Supplementary Fig. 9c), smoking in bladder and lung (Supplementary Fig. 9d,f), and isocitrate dehydrogenase-1 *(1DH1)* mutational status in brain cancers (Fig. 4g). Each of these findings are in agreement with the current knowledge about determinants of DNA methylation^15, 46–48^. These results further validate our models and also emphasize the importance of the contribution observed for the SGOC variables (including the met cycle).

Previous work has shown that expression of enzymes across different regions of the SGOC network is predictive of metabolic flux through the network^24^. Notably, we observed that several SGOC genes are consistently among the highly ranked variables by both Random Forest and Elastic Net models in multiple cancer types. Therefore, to understand which features of SGOC metabolism contribute to the interaction with methylation, we defined a sub-network that was commonly highly ranked by the models in multiple cancer types (Fig. 4h; Online Methods). This SGOC sub-network comprises the MAT enzymes in the met cycle (MAT2B and MAT2A), as well as enzymes within serine-glycine metabolism such as phosphoglycerate dehydrogenase (PHGDH) and glycine amidinotransferase (GATM) (Fig. 4h). We generally observed negative associations between DNA methylation and expression of PHGDH and GATM, but positive associations with expression of MAT enzymes. A cautionary note however is that in many disease states, levels of particular metabolites in the methionine cycle substantially deviate from physiological ranges, thus activating compensatory mechanism and leading to correlation with DNA methylation in directions opposite of what would be expected from the biochemistry of the reactions^49^. Therefore, when interpreting the direction of correlations between metabolic enzyme levels and DNA methylation, it is important to note that they not only depend on the stoichiometry of the corresponding enzymatic reactions, but also on endogenous abundance of the related metabolites. Together, our results suggest that a particular flux configuration through the SGOC metabolic network— which previous studies have shown to be predictable from gene expression patterns^24^—may be important for regulation of DNA methylation.

### Cancer pathogenesis of metabolically regulated DNA methylation

Involvement of the met cycle in promoter and gene body methylation at cancer genes suggests a potential implication for this metabolic pathway in explaining part of the variability in cancer pathogenesis and patient outcome. To further assess this relationship, we divided patients in each cancer type into two groups based on overall predictability of their cancer loci methylation by the met cycle (see Online Methods). We then compared survival rates between the two groups (“predictable” by met cycle vs. “not predictable” by met cycle) in each cancer type using the Kaplan-Meier estimator^50^ (Fig. 5a–h). An improved overall survival for the “predictable” group was observed, although the magnitude of this trend varied depending on cancer type with brain, kidney, liver, and colon cancers showing statistically significant differences (log-rank test p-values: brain = 3.92e-05, liver = 0.0048, kidney = 0.0085, colon = 0.04) (Fig. 5a–d). The difference in survival between the predictable and non-predictable groups was not significant in the rest of the cancers studied here (Fig. 5e–h), possibly explained by limited power due to data censoring at later time points. The overall patterns however suggest that the regulation of DNA methylation by the met cycle may be important in maintaining a normal epigenome, and disruption of this relationship in specific subtypes of tumors can lead to high epigenetic stochasticity in those tumors that correspond to poor clinical outcomes. This is consistent with a previous study that showed DNA methylation stochasticity increased across samples with increasing malignancy (from normal to adenoma to carcinoma)^6^.

**Figure 5.**
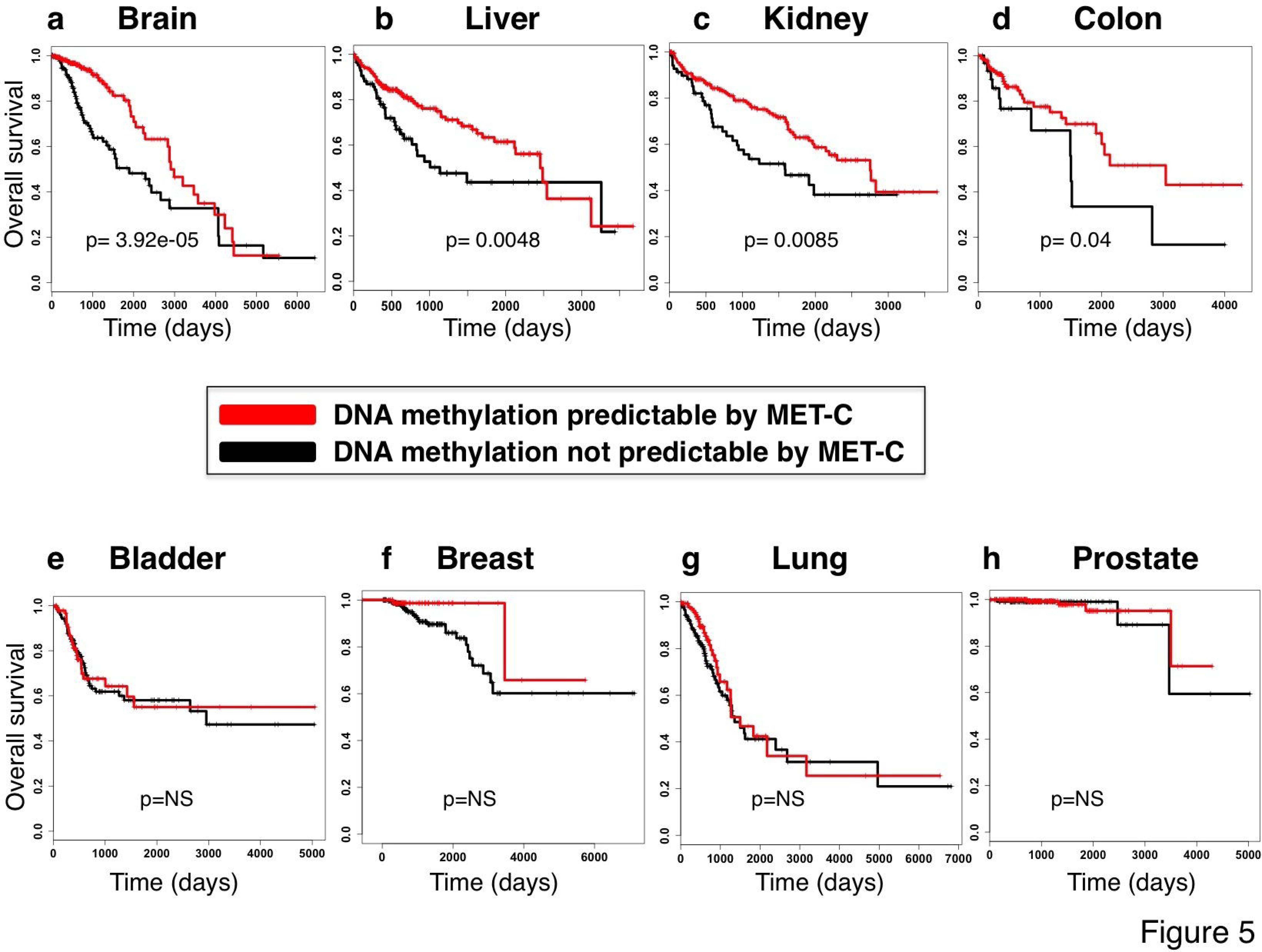
Implication of metabolic regulation of methylation in patient survival.

a-d) Kaplan-Meier curves are depicted comparing groups of patients wherein cancer gene methylation was predictable (red) or not predictable (black) by the met cycle variables (see Online Methods). Overall survival in days is plotted in each case and censored subjects are shown by vertical tick marks (Online Methods). Log-rank test p-value between the two groups is reported. Survival analysis results and log-rank test p-values are shown for brain, liver, kidney, and colon cancers respectively.

e-h) Survival analysis results as described above are reported for bladder, breast, lung, and prostate cancers, respectively. Log-rank test p-values showed no significant difference between the “predictable” and “not predictable” groups (“NS” = not significant) (Sample sizes: breast = 770, lung = 450, liver = 374, brain = 534, bladder = 408, kidney = 316, prostate = 424, colon = 198)

To validate the results of our survival analyses, we applied multivariate cox regression models to account for covariates such as mutations and clinical factors that are know to be associated with survival rates (Online Methods). We performed this test in the cases of brain, liver, and kidney cancers were the univariate analyses found highly significant differences between the predictable and non-predictable groups (Fig. 5a–c). The models including covariates still showed a significant difference (p < 0.05) between the predictable and non-predictable groups of patients even after taking mutational and clinical factors into account (see Online Methods for the list of covariates considered in each cancer), suggesting that a unique part of variation in survival may be explained by epigenetic regulation. We next tried to further validate our results through comparison with independent analyses of the TCGA data by the cBioPortal for Cancer Genomics (cBioPortal)^51^ and Prediction of Clinical Outcome from Genomic profiles (PRECOG)^52^. These analyses found lower survival in prostate cancer patients harboring tumors with deep deletions in the met cycle genes (Supplementary Fig. 10a), and higher survival in kidney cancer patients where the met cycle enzymes are over-expressed (Supplementary Fig. 10b), respectively. These results confirm a relationship between met cycle and survival in the same direction as predicted by our hypothesis.

## Discussion

In this study, we conducted a pan-cancer TCGA analysis of the molecular and clinical contributions to within-cancer (inter-individual) variation in DNA methylation. Through several lines of integrated analysis, we found the overall expression of both the methionine cycle and SGOC network to be strong predictors of multiple aspects of DNA methylation and consistently ranked as one of the highest contributing factors to cancer-associated DNA methylation such as methylation of numerous cancer genes. Within the methionine cycle, we consistently observed a more significant contribution from MAT2B and BHMT2, suggesting that the regulation may be occurring at these enzymatic steps. MAT2B is the enzyme that converts methionine to SAM, therefore it is expected that this enzyme affects SAM levels more directly than other metabolic enzymes. The significance of BHMT2 but not MTR suggests that metabolism of choline and betaine may be more prevalent than folates in cases where one-carbon metabolism fuels DNA methylation.

We introduced a novel approach to identify chromatin regions with strong correlations between DNA methylation and metabolic enzyme levels. The identified regions for the met cycle enzymes significantly overlapped with histone modifications, consistent with enzymatic cross-talk between the two epigenetic processes^35^. The enrichment of gene signatures of repressing histone marks such as H3K27me3 in all cancers points to a possible role for the met cycle in maintenance of DNA methylation at silenced loci. Previous studies have reported aberrant methylation of transcriptionally repressed genes in cancer^53^. In fact, heterochromatin instability arising from increased variability in DNA methylation is a phenomenon observed in many cancers and is thought to contribute to epigenetic plasticity and tumor progression^4, 54^. Our results provide evidence for this model of dysregulated cancer epigenome and further suggest that disruption of the regulation of DNA methylation by the met cycle—which can be a cause or consequence of tumorigenesis— may be one of the sources of methylation stochasticity leading to higher malignancy. Survival analyses confirm that tumors with a weaker association between their cancer gene methylation and the met cycle expression are more malignant in comparison to tumors wherein this relationship is closer to normal. In addition to epigenetic overlaps, genes with important tissue-specific functions and disease states were also found to fall under the metabolism-DNA methylation peaks. DNA methylation at cell-type related disease and lineage-specific genes has previously been shown to be dynamic and functionally important^14^. Our results further strengthen the idea that met cycle regulation of methylation is strongly associated with normal tissue function.

Application of the integrative modeling to cancer genes revealed a major role for MAT enzymes (MAT2B and MAT2A), as well as PHGDH and GATM— enzymes involved in serine and glycine metabolism, respectively. Importantly, MAT2B and MAT2A have been shown to co-localize in nuclei and bind DNA through complex formation with chromatin binding proteins providing direct evidence for the role of these enzymes in regulation of transcription via methylation^55^. PHGDH diverts the glycolytic flux into the de novo serine synthesis pathway that allows glycolysis to provide methyl units. GATM diverts glycine into the creatine synthesis pathway in which SAM is consumed to produce creatine^56^. Creatine synthesis is therefore in competition with the methionine cycle over cellular pools of SAM, explaining why enzymes within the serine-glycine metabolism generally tend to be negatively correlated with the met cycle and DNA methylation.

Overall, this study provides the first comprehensive quantification of the determinants of inter-individual DNA methylation variation in human cancers. The activity of the methionine cycle that emerges in these findings could be either sensed directly by the DNA, or indirectly through interplay with dynamic histone methylation, which itself is tightly regulated by the status of methionine metabolism^21^. Due to limitation in the coverage of the DNA methylation arrays, it remains to be determined if our findings are generalizable to methylation across the entire genome including all non-CpG methylation sites as well as hydroxy-methylation sites. Nevertheless these findings altogether identify metabolism as a major determinant of DNA methylation status in human cancer. It is important to note that the current TCGA dataset contains one sample per individual tumor and therefore our conclusions do not necessarily explain the variation in clonal populations within a given tumor. Future studies using multiple samples per tumor or single cell epigenomics are therefore required to characterize the determinants of intra-tumor epigenetic heterogeneity. Finally, our study identifies a role for altered tumor metabolism in explaining DNA methylation, while the sources of alterations in metabolism itself remain to be elucidated but can be addressed using similar approaches.

## Acknowledgments

We thank Sam Mentch for useful comments and help with the graphics. This work was supported by grants R00CA168997 and R01CA193256 to J.W.L. from the National Institutes of Health. M.M. was also supported by T32GM007617 training grant from the NIH and a Graduate Fellowship from the Duke University School of Medicine.

## Online Methods

### Data curation

Publically available genome-wide mRNA expression and DNA methylation data were downloaded from the cancer genome atlas (TCGA) portal(*https://tcga-data.nci.nih.gov/tcga/*). In order to increase consistency and minimize unwanted variations, only samples processed using RNASEQ-V2 with level-3 gene-normalized RNA-seq by Expectation Maximization (RSEM) values for gene expression, and level-3 beta-values from Illumina Infinium HumanMethylation450K BeadChip data for DNA methylation were included in the study. We selected the following 8 cancer types wherein the number of available samples analyzed on both platforms was sufficiently large for machine-learning calculations: 770 samples of breast invasive carcinoma (BRCA), 450 samples of lung adenocarcinoma (LUAD), 374 samples of liver hepatocellular carcinoma (LIHC), 534 samples of brain lower grade glioma (LGG), 408 samples of bladder urothelial carcinoma (BLCA), 316 samples of kidney renal clear cell carcinoma (KIRC), 424 samples of prostate adenocarcinoma (PRAD), and 198 samples of colon adenocarcinoma (COAD). Somatic mutations with a frequency of 5% or higher, and Genomic Identification of Significant Targets in Cancer (GISTIC) values for copy number alterations with a frequency of 15% or higher according to the cBioPortal^51^ were obtained and included in the models. Clinical and follow-up data were downloaded via the TCGA-Assembler^57^.

### Assessment of batch effects

We used the TCGA Batch Effects online tool(http://bioinformatics.mdanderson.org/tcgabatcheffects) to check for the existence of batch effects in the data used in our study. For each cancer types in our study, both the DNA methylation and the RNA-seq batch effects were negligible (Dispersion Separability Criterion (DSC) score < 0.5 for all sample batches included in the study).

### DNA methylation

The Illumina Infinium Human Methylation450K BeadChip consists of more than 450,000 probes across the genome covering CpG sites within and outside of CpG islands as well as non-CpG methylation sites identified in embryonic stem cells (*see:http://www.illumina.com/products/methylation_450_beadchip_kits.html*). We first filtered all probes with more than 80% missing values across each cancer type. Global DNA methylation was then defined as the average beta-value across all remaining probes for each sample (Supplementary Fig. 1a). Sex chromosomes were also excluded from all subsequent analyses of DNA methylation. In order to assess local DNA methylation, we divided the genome into 10kb intervals and calculated the average beta value across all probes within each bin. We then filtered regions where variation in methylation was modest (standard deviation < 0.2 across each dataset). The average beta-value across all remaining 10kb regions was then calculated for each sample individually and plotted in Supplementary Fig. 1c. In order to study DNA methylation at cancer loci, probes that mapped to each gene according to Illumina annotations were identified. Promoter DNA methylation was then defined as the average beta value across all probes mapping to a given gene and falling within one of the following positional categories based on Illumina chip annotation information: “TSS1500”, “TSS200”, or “5’UTR”. Gene body methylation for each gene was defined as the average beta value across all probes mapping to a given gene and falling in “1^st^ exon”, “Body”, or “3’UTR” based on the annotation. Promoter and gene body methylation were separately modeled for each of the cancer genes in the study (Fig. 4).

### Gene expression

Log-transformed gene normalized RSEM values were used as expression levels and low-expression genes in each dataset were defined as having less than 70% of the samples with a count value larger than 3. Such genes were removed from further analysis.

### Gene expression variables included in the integrative models

In addition to the major enzymes in the met cycle (MAT2B, MTR, BHMT2, AHCY), four classes of expression variables with potential links to DNA methylation were also included in the integrative models (see Supplementary Table 1 for the complete list of variables). The four classes are described in the following:

“Other SGOC enzymes”: Serine, glycine, one-carbon (SGOC) metabolic genes from our previous network reconstruction were included ^24^. In order to separately assess the effect of the met cycle from the rest of the network, we excluded the met cycle enzymes from this class and treated them as a separate class (“Methionine cycle enzymes”).

“Chromatin Remodelers”: A list of human chromatin remodelers and DNA methylation machinery was constructed by combining the Gene Ontology (GO) chromatin modifiers list, GO chromatin remodelers list^58^, and methylated DNA binding proteins and demethylases^59^.

“Transcription factors”: For each cancer type, transcription factors important in the pathogenesis or subtype specification based on previous literature were included^60^. “SAM-metabolizing enzymes”: DNA methyltransferases and other SAM-consuming enzymes (except for MAT enzymes already included in the class “Methionine cycle enzymes”) according to Human Cyc^61^ were included in this class.

### Mutations included in the integrative models

For each cancer type, genes with frequent somatic mutations (minimum frequency of 5%) among the TCGA cohort according to the cBioPortal^51^ summary table (TCGA, Provisional) were obtained. The transposed matrix of individual barcodes and mutations in the selected genes was downloaded from the cBioPortal for each of the 8 cancers in this study. See Supplementary Table 1 for a complete list of somatic mutations considered in each cancer type.

### Copy number alterations included in the integrative models

For each cancer type, genes with frequent copy number alterations (minimum frequency of 15%) among the TCGA cohort according to the cBioPortal^51^ summary table (TCGA, Provisional) were obtained. The transposed matrix of individual barcodes and putative copy number alteration calls by GISTIC^62^ for the selected genes was downloaded from the cBioPortal for each of the 8 cancers in this study (Values of putative copy number calls determined using GISTIC 2.0: −2 = homozygous deletion; −1 = hemizygous deletion; 0 = neutral / no change; 1 = gain; 2 = high-level amplification). See Supplementary Table 1 for a complete list of copy number alterations considered in each cancer type.

### Clinical factors included in the integrative models

For each cancer type, clinical information was downloaded through the TCGA-Assembler^57^. All clinical attributes were included for each cancer type with the exception of the ones filtered out due to missing data for all samples or factors with the same level across all samples. See Supplementary Table 1 for a complete list of clinical attributes considered in each cancer type.

### Predictive modeling and variable ranking using the Random Forest algorithm

The Random Forest is a machine-learning algorithm that generates predictions by averaging over a collection of randomized decision trees. Since successive trees are built with bootstrap samples, the algorithm is robust to over-fitting, and also those samples that are left out (the *out-of-bag* (OOB) samples) can be used to quantify the contribution that prediction variables make to the overall response. The Random Forest method is designed to accommodate nonlinearities between the response and prediction variables as well as unknown interactions among the variables^29,63,64^. We used the R package “randomForest”^65^ and performed 3-fold cross validation by manually dividing the samples in each cancer type into 3 training and test subsets. To build each forest, tree size was set to 500 and the “importance” parameter was set to “TRUE” in the R function “randomForest” so as to provide estimates for the importance of prediction variables.

Missing data were imputed using the “na.roughfix” function in the “randomForest” package. We obtained separate measures of importance for each variable from each Random Forest run. These importance scores are calculated as the percent increase in the mean squared prediction error on the OOB samples when a given variable is permutated. Variables were ranked based on average importance scores across all cross validation folds. Prediction errors were calculated as the mean squared difference between the predicted vs. the observed methylation values for the test set samples. The square root of the mean squared error (MSE) has the same scale as the response (DNA methylation beta values in this case), and is therefore a direct measure of the accuracy with which predictions were made. (Fig. 1c; Supplementary Fig. 6a,b).

### Predictive modeling and variable selection using the Elastic Net algorithm

Elastic Net is a penalized regression approach for variable selection and quantitative inference that identifies linear combinations of unique variables that contribute to a response variable such as the amount of DNA methylation. The algorithm was developed and benchmarked to avoid over-fitting in statistical modeling of high-dimensional data containing collinearity^27^. We applied the Elastic Net algorithm using the R package “glmnet”^66^. Elastic Net performs variable selection by minimizing a regularized cost function using the following equation

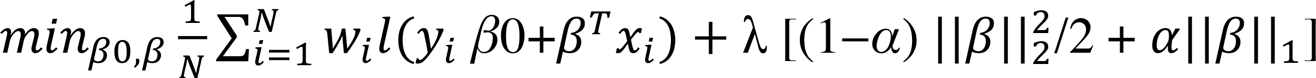

where lambda is the tuning parameter and alpha is the Elastic Net penalty term. For each cancer type, the samples were divided into 3 independent test subsets (3-fold cross validation), and separate models were generated using each training subset. Using a grid of different tuning parameter values, we found the lambda that minimized the mean squared error using 5-fold cross validation within each training set for each model separately. The value of alpha was set to α = 0.5 to handle potential correlated variables. Finally, for each variable, average coefficient across the 3 independent models was calculated for each region and each cancer type. Due to the existence of categorical factors among our variables (for which scaling is not appropriate), we also calculated the selection rate as an alternative measure of variable importance referred to as “variable usage” in the manuscript. Variable usage was measured as the fraction of times across all cross validation folds that a variable was selected by the Elastic Net to be included in the final model (Supplementary Fig. 6c; Fig.4 f,g; Supplementary Fig. 9a-f). Finally, prediction errors were calculated as the squared difference (mean squared error (MSE)) between the predicted and measured DNA methylation values for the test sets (Fig. 1c; Supplementary Fig. 6a,b).

### Variable class contributions to DNA methylation

Variables were functionally categorized into the following 8 classes: “Methionine cycle enzymes”, “Other SGOC enzymes”, “Chromatin remodeling factors”, “Transcription factors”, “SAM metabolizing enzymes”, “Clinical factors”, “Copy number variations”, and “Mutations”. Results of the integrative modeling were summarized and reported in terms of the average contribution from each of the above functional classes in explaining DNA methylation variation. Variable importance scores from Random Forest models were averaged across all variables within a given class, and an overall class importance score was calculated. In the case of Elastic Net models, variable usage as described in the previous section, was averaged across variables in each class and an average percentage showing selection rate was calculated. Finally, classes were ranked in each cancer type according to their average contribution and the overall class ranks were plotted in Fig. 2a, Supplementary Fig. 2c,d, and Supplementary Fig. 6d,e.

### Comparison to variance-matched random gene expression controls

A set of 100 randomly selected genes from the genome with similar cross-sample variation in expression as our original gene expression variables (TFs, SGOC, MET-C, SAM, and RMs) were considered. We performed this test on local DNA methylations (all variable 10kb regions) in brain cancer (LGG) as an example and repeated the integrative modeling using this set of randomly selected genes in addition to all other variables present in the original models. All gene expression variables were then ranked using a similar approach as described above. To compare our original gene expression variables with the variance-matched random genes, the ranks across all models were averaged (Fig. 1d), and p-values were obtained from one-tailed Mann-Whitney non-parametric test between the two groups from Elastic Net and Random Forest.

### Distance to nearest gene transcriptional start site (TSS)

Selected 10kb regions were converted to genomic range objects using the R package “GenomicRanges”^67^. The distance to single nearest gene's transcription start site (TSS) was found using Genomic Regions Enrichment of Annotations Tools (GREAT)^68^.

Genomic regions are associated with nearby genes by first assigning a regulatory domain to every gene in the genome, and then finding genes whose regulatory domains overlap with a given genomic region. We set the association rule parameter in GREAT to “Single nearest gene” with a maximum extension of 1000kb for definition of regulatory domains. Density plots of distance to TSS are depicted in Supplementary Fig. 2b. The same approach was used for annotating peaks obtained from Fig. 3 (density plots shown in Supplementary Fig. 3a,c). To obtain the distribution of Illumina probe densities around the TSS, we randomly selected 10000 probes across the arrays and applied the above-described approach to measure the distance to nearest gene's TSS for each probe. Density plots were obtained for the purpose of comparison with the distribution of metabolically regulated peaks (Supplementary Fig. 3b,d).

### Identification of metabolically regulated genomic regions

To find peaks of strong association between the met cycle and DNA methylation, we designed a novel scanning method by applying the idea of Manhattan plots from e-QTL analyses^69^ to DNA methylation data. In each cancer type, we first selected one of the major enzymes in the met cycle with the highest overall Spearman correlation with global and local DNA methylations (*BHMT2* in brain, breast, prostate, and liver; *MAT2B* in lung and bladder; and *AHCY* in colon and kidney cancers), and calculated the Spearman correlation between its expression and the beta value of each individual probe across the genome. We then sorted the probes according to genomic coordinates and aligned the – log10 of the p-values obtained from the Spearman correlations along the chromosomes. Next, we applied a sliding window scan for regions of strong association across the genome separately in each cancer type (Fig. 3a). For this, probes with the highest correlations (top 10% across the genome) were located and a 6kb window (+3kb and −3kb) flanking the genomic coordinate of the original probe was scanned. A region was reported as a “peak” if the following criteria were met: 1-Region included at least 3 probes with a correlation in the same direction as the original probe (positive or negative); 2-At least 80% of all probes within the region had a significant (p < 0.00001) correlations with met cycle expression. After applying these filters, the selected regions were annotated and genes overlapping with each of the peaks were used for subsequent pathway enrichment analyses. Given the window size and the above criteria, the majority of the identified peaks only overlapped with one unique gene (see Supplementary Table 2 for a complete list of all identified peaks).

To assess potential bias toward highly methylated regions in the identified regions where correlation of methylation with met cycle expression peaks, we tested 2000 randomly selected probes across the genome. We then evaluated the association between methylation of each probe with the value of its Spearman correlation rho with met cycle expression— we used AHCY in colon cancer as an example in this test (Fig. 3b).

Finally, an additional filter was applied to rank the identified peaks according to peak shape. For this, the aligned correlation coefficients in each region were assessed with respect to whether they formed a peak according to an information theory score calculated by the R function “turnpoints” (refer to R package “pastecs”^70^). This function finds all turning points (peaks and pits) in a series of points (in this case, aligned correlation coefficients), and calculates the information quantity of each turning point using Kendall's information theory. Finally, it measures a p-value against a random distribution of the turning points in a given series, with smaller p-values corresponding to less random shape and a higher probability of a turning point corresponding to a real peak or pit. We selected regions containing turning points with the most significant p-values (lowest 20%) in each cancer type and subsequently tested them for specificity for the met cycle as described in the following section.

### Test of specificity of peaks for the met cycle

Each of the selected peaks was tested for specificity of their correlations with the met cycle expression (vs. gene expression in general). For this, 500 genes were randomly selected from the genome in each cancer type, and the Spearman correlation coefficient was measured between their expression and the methylation of every probe within a given peak. The fraction of significant correlations was calculated for all of the 500 genes as well as for the met cycle gene. A randomization q-value was calculated for the met cycle gene by comparing it to the distribution of the correlations calculated for the 500 random genes. This procedure was repeated separately for each peak in each cancer type and the results are summarized in Supplementary Fig. 6a,b.

### Pathway enrichment analyses

Peaks were annotated according to Illumina information and UCSC Ref gene names for genes overlapping with the identified peaks were extracted. Pathway enrichment analysis was performed on the resulting gene list for each cancer type using Enrichr^32^. Combined scores from Enrichr were used to rank pathways. The Combined score “c” is defined as *c = log(p)*z* where p refers to the p-value from the Fisher's exact test and z is the z-score indicating the deviation from the expected rank. Enrichr first calculates Fisher's exact p-values for many random gene sets to generate a distribution of expected p-values for each pathway in their pathway library. The z-score for deviation from this expected rank is therefore an alternative ranking score and the combined score is considered a corrected form of the enrichment score and p-value, which we used to sort pathways in Fig. 3c–f and Supplementary Fig. 5a-d. All gene sets in Fig. 3c–f and Supplementary Fig. 5 had Fisher's exact p-values < 0.05, and the most highly enriched sets are shown ranked by the combined enrichment scores. Gene set names used in Fig. 3c–f and Supplementary Fig. 5 follow the convention used and described by Enrichr (*http://amp.pharm.mssm.edu/Enrichr/#stats*). Briefly, gene ontology (GO) sets are shown by GO numbers in parenthesis following their name, epigenetic modifications from the ENCODE histone modifications 2015 project are shown by “-hg19” following gene set names to be distinguished from those from the Epigenomics Roadmap project, gene sets from the Cancer Cell line Encyclopedia are shown by cell line names following cancer type in upper case, disease signatures from the gene expression omnibus (GEO) are shown in upper case followed by GSE accession numbers, KEGG 2015 and the Human Gene Atlas gene sets are shown in lower case. Refer to Enrichr for a complete list of all gene sets included in more than 70 libraries.

### Cancer genes

A list of 12 cancer drivers common in multiple human cancers was considered^71^ (tumor protein p53 (*TP53*), phosphate and tensin homolog (*PTEN*), neuroblastoma RAS viral oncogene homolog (*NRAS*), epidermal growth factor receptor (*EGFR*), isocitrate dehydrogenase 1(*aIDH1*), isocitrate dehydrogenase 2 (*IDH2*), CCCTC-binding factor (*CTCF*), von Hippel-Lindau tumor suppressor, E3 ubiquitin protein ligase (*VHL*), catenin beta 1 (*CTNNB1*), nuclear factor erythroid-2 like 2 (*NFE2L2*), phosphoinositide-3-kinase, regulatory subunit 1 (*PIK3R1*), and ms-related tyrosine kinase 3 (*FLT3*)). These genes were consistently identified as candidate cancer drivers by 4 independent positive selection detection algorithms in a comprehensive pan-cancer analysis of thousands of TCGA tumors^71^. We added to this list, well-known cancer drivers not included in the above list (Kirsten rat sarcoma viral oncogene homolog (*KRAS*), B-Raf proto-oncogene, serine/threonine kinase (*BRAF*), phosphatidylinositol-4,5-bisphosphate 3-kinase, catalytic subunit alpha (*PIK3CA*), and breast cancer 1, early onset (*BRCA1*)). In addition to these common cancer drivers, we also considered a number of cancer type-specific genes including receptors important in specific subtypes of cancers (estrogen receptor 1(ESR1), androgen receptor (AR), erb-b2 receptor tyrosine kinase 2 (*ERBB2*)). Finally, cancer genes frequently aberrantly methylated in human cancers were also considered^42^ (RAS association domain family member-1 (*RASSF1*), glutathione S-transferase pi 1 (*GSTP1*), adenomatous polyposis coli (*APC*), and O-6-methylguanine-DNA methyltransferase (*MGMT*)), together constructing a list of 23 cancer genes for detailed analysis of DNA methylation shown in Fig. 4.

### Evaluation of modeling performance using randomized responses

In order to test the reliability of the variable contribution results obtained from our gene-specific DNA methylation models, we built two different randomized data sets as control responses, each with the same dimensions as the original response dataset (i.e. the cancer gene DNA methylations). In the first case, we permuted the DNA methylation values of each cancer gene, and repeated the modeling using the met cycle variables. In the second case, we generated random beta-values (from uniform distribution in the range of 0-1) and used those as the response variables in the calculations. We then compared average met cycle variable importance (Random Forest) and variable usage (Elastic Net) from prediction of true cancer gene methylations vs. permuted methylations and randomly generated responses. The Kolmogrov-Smirnov test p-values were calculated between the distributions as illustrated in Supplementary Fig. 7b.

### Evaluation of modeling performance using randomized prediction variables

Using prostate cancer as an example, we performed simulation tests to determine whether the Random Forest as a methodology is able to utilize the information in the prediction variables beyond what could be expected if the predictors were only random noise and unrelated to the response. To investigate this, we modeled methylation in the prostate cancer dataset at 3 example cancer loci (*GSTP1*, *RASSF1*, *and PITX2*). These genes were selected based on previous evidence indicating the critical importance of their aberrant methylation in prostate cancer^42, 72^. As controls, we generated 3 additional datasets. For the first dataset, we copied the exact response as the *GSTP1* methylation, but randomly generated a predictor variable set of the same dimensions as the original variable set by sampling from a standard normal distribution. That is, each observation on each variable is a sample from a normal distribution of unit variance and should therefore have no relationship to the response. The other two datasets were generated in the same fashion, using *RASSF1* and *PITX2* methylation as responses and randomly generated variable sets as predictors. To assess the performance of the Random Forest computations, we compared the mean squared error (MSE) from predictions made using the original data with those made by the datasets consisting of random variables unrelated to the responses. For each of the three responses, we randomly divided the data into training and test sets, generated a total of 100 simulations consisting of 500 decision trees, and compared the resulting MSEs of the predictions made on the test points. Results are summarized in Supplementary Fig. 8a-d.

To quantify the improvement in the Random Forest algorithm by using the original variables over the randomly simulated variables, we defined an improvement metric (MSE-Imp), describing the relative improvement in prediction accuracy:

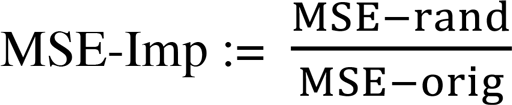

where MSE-rand is the average MSE calculated using the random simulated variables and MSE-orig is the average MSE calculated using the original variables.

In this test, another simulated dataset of the same dimensions as the original dataset was generated where the variables and response were linearly related via the following equation:

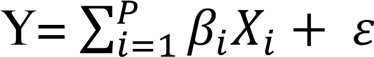

To generate this linear dataset, we sampled the value of each prediction variable Xi from a standard normal distribution and the noise *ε* from a normal distribution with mean 0 and standard deviation 0.05. The values of the coefficients *βi* were selected uniformly at random from the interval [0,1]. We then measured the MSE improvement (MSE-Imp) for the linear dataset using the same approach as MSE improvements for the original datasets were calculated (explained in the previous paragraph). This allowed us to compare a linearly simulated dataset to our real dataset. Results are shown in Supplementary Fig. 8e-f.

### Network construction

Genes in the serine, glycine, one-carbon (SGOC) network (including the met cycle genes) that were among the most highly ranked variables (top 15%) in at least 4 of the cancer datasets according to the Elastic Net models and at least 3 of the cancer datasets according to the Random Forest models were selected. A metabolic network consisting of these enzymes was then constructed using MetScape^73^ where nodes represent genes and metabolites, and edges represent biochemical links. We fixed the node size for metabolites but adjusted node sizes for genes to correspond to the number of cancers in which each variable was highly ranked (among the top 15% of all variables) (Fig. 4g). For nodes not directly connected to the rest of the network, we manually added dashed lines where appropriate.

### Survival analyses

In each cancer type, the average error of prediction of DNA methylation at cancer loci was measured for each patient across all Elastic Net models using only met cycle variables for prediction. Patients were then divided into 2 groups based on predictability of their methylation by the met cycle activity (“predictable” = below-median prediction error, “not predictable” = above-median prediction error). To estimate overall survival time, “days-to-death” was used with vital status information and last follow-up date used to right-censor subjects (subjects alive at last follow-up were censored from the analysis beyond their last follow up date). The relationship between survival and predictability was then analyzed using the “survfit” function in the R package “survival”^74^ and visualized by Kaplan-Meier curves. Log rank test p-values were calculated by fitting models of overall survival to patients' “predictability” group assignments using the “survdiff” function in the survival package for each cancer type separately. Results are depicted in Fig. 5.

### Multivariate cox regression

In the three cancer types (brain, liver, and kidney) where univariate analysis showed a highly significant difference in survival between the predictable and non-predictable groups as described above, and also the sample size allowed for sufficient power to perform multivariate analysis, we used relevant clinical and mutational factors as covariates and repeated the survival analysis. The following factors were individually tested as covariates in separate models of overall survival along with “predictability” status as the fixed effect: Brian cancer: all frequent somatic mutations (see Supplementary Table 1 for the complete list), histological diagnosis, age, gender, and initial weight; Liver cancer: all frequent somatic mutations (see Supplementary Table 1 for the complete list), tumor stage, history of other malignancies, and residual tumor; Kidney cancer: all frequent somatic mutations (see Supplementary Table 1 for the complete list), age, and race. In each case, the results of regression using the “coxph()” function in R provided the p-value for the significance of the predictability status when modeling overall survival in the presence of covariates.

### Software

^*^All computational and statistical analyses were done using R 3.1.2 ^75^. Distribution plots, box-plots, scatter-plots, and bar-plots were made in GraphPad Prism version 6 (GraphPad Software, San Diego California USA, *www.graphpad.com*). Circular plots were generated using Circos^76^.

### Code availability

R script is available through the following Github repository (https://github.com/mahyam/DNA-methylation-and-metabolism-R-code).

## References

1. Jones, P.A. Functions of DNA methylation: islands, start sites, gene bodies and beyond. Nature reviews. Genetics 13, 484–492 (2012).

2. Schubeler, D. Function and information content of DNA methylation. Nature 517, 321–326 (2015).

3. Hamidi, T., Singh, A.K. & Chen, T. Genetic alterations of DNA methylation machinery in human diseases. Epigenomics 7, 247–265 (2015).

4. Hansen, K.D., et al. Increased methylation variation in epigenetic domains across cancer types. Nature genetics 43, 768–775 (2011).

5. Mack, S.C., et al. Epigenomic alterations define lethal CIMP-positive ependymomas of infancy. Nature 506, 445–450 (2014).

6. Timp, W. & Feinberg, A.P. Cancer as a dysregulated epigenome allowing cellular growth advantage at the expense of the host. Nature reviews. Cancer 13, 497–510(2013).

7. Witte, T., Plass, C. & Gerhauser, C. Pan-cancer patterns of DNA methylation. Genome medicine 6, 66 (2014).

8. Ehrlich, M. & Lacey, M. DNA hypomethylation and hemimethylation in cancer. Advances in experimental medicine and biology 754, 31–56 (2013).

9. Jones, P.A. & Baylin, S.B. The epigenomics of cancer. Cell 128, 683–692 (2007).

10. Lopez-Serra, P. & Esteller, M. DNA methylation-associated silencing of tumor-suppressor microRNAs in cancer. Oncogene 31, 1609–1622 (2012).

11. Berman, B.P., et al. Regions of focal DNA hypermethylation and long-range hypomethylation in colorectal cancer coincide with nuclear lamina-associated domains. Nature genetics 44, 40–46 (2012).

12. Gaidatzis, D., et al. DNA sequence explains seemingly disordered methylation levels in partially methylated domains of Mammalian genomes. PLoS genetics 10, e1004143 (2014).

13. Lokk, K., et al. DNA methylome profiling of human tissues identifies global and tissue-specific methylation patterns. Genome biology 15, r54 (2014).

14. Ziller, M.J., et al. Charting a dynamic DNA methylation landscape of the human genome. Nature 500, 477–481 (2013).

15. van Dongen, J., et al. Genetic and environmental influences interact with age and sex in shaping the human methylome. Nature communications 7, 11115 (2016).

16. Locasale, J.W. Serine, glycine and one-carbon units: cancer metabolism in full circle. Nature reviews. Cancer 13, 572–583 (2013).

17. Gut, P. & Verdin, E. The nexus of chromatin regulation and intermediary metabolism. Nature 502, 489–498 (2013).

18. Kaelin, W.G., Jr. & McKnight, S.L. Influence of metabolism on epigenetics and disease. Cell 153, 56–69 (2013).

19. Sahar, S. & Sassone-Corsi, P. Metabolism and cancer: the circadian clock connection. Nature reviews. Cancer 9, 886–896 (2009).

20. Anderson, O.S., Sant, K.E. & Dolinoy, D.C. Nutrition and epigenetics: an interplay of dietary methyl donors, one-carbon metabolism and DNA methylation. The Journal of nutritional biochemistry 23, 853–859 (2012).

21. Mentch, S.J., et al. Histone Methylation Dynamics and Gene Regulation Occur through the Sensing of One-Carbon Metabolism. Cell metabolism 22, 861–873 (2015).

22. Pfalzer, A.C., et al. S-adenosylmethionine mediates inhibition of inflammatory response and changes in DNA methylation in human macrophages. Physiological genomics 46, 617–623 (2014).

23. Cancer Genome Atlas Research, N., et al. The Cancer Genome Atlas Pan-Cancer analysis project. Nature genetics 45, 1113–1120 (2013).

24. Mehrmohamadi, M., Liu, X., Shestov, A.A. & Locasale, J.W. Characterization of the usage of the serine metabolic network in human cancer. Cell reports 9, 1507–1519 (2014).

25. Duncan, C.G., et al. A heterozygous IDH1R132H/WT mutation induces genome-wide alterations in DNA methylation. Genome research 22, 2339–2355 (2012).

26. Zou, H. & Hastie, T. Regularization and variable selection via the Elastic Net. Journal of the Royal Statistical Society Series B, 301–320 (2005).

27. Waldmann, P., Meszaros, G., Gredler, B., Fuerst, C. & Solkner, J. Evaluation of the lasso and the elastic net in genome-wide association studies. Frontiers in genetics 4, 270 (2013).

28. Breiman, L. Random Forests. Machine Learning 45, 5–32 (2001).

29. Chen, X. & Ishwaran, H. Random forests for genomic data analysis. Genomics 99, 323–329 (2012).

30. Stadler, M.B., et al. DNA-binding factors shape the mouse methylome at distal regulatory regions. Nature 480, 490–495 (2011).

31. Feldmann, A., et al. Transcription factor occupancy can mediate active turnover of DNA methylation at regulatory regions. PLoS genetics 9, e1003994 (2013).

32. Chen, E.Y., et al. Enrichr: interactive and collaborative HTML5 gene list enrichment analysis tool. BMC bioinformatics 14, 128 (2013).

33. Consortium, E.P. An integrated encyclopedia of DNA elements in the human genome. Nature 489, 57–74 (2012).

34. Roadmap Epigenomics, C., et al. Integrative analysis of 111 reference human epigenomes. Nature 518, 317–330 (2015).

35. Cedar, H. & Bergman, Y. Linking DNA methylation and histone modification: patterns and paradigms. Nature reviews. Genetics 10, 295–304 (2009).

36. Hon, G.C., et al. Global DNA hypomethylation coupled to repressive chromatin domain formation and gene silencing in breast cancer. Genome research 22, 246–258 (2012).

37. Ho, V., Ashbury, J.E., Taylor, S., Vanner, S. & King, W.D. Gene-specific DNA methylation of DNMT3B and MTHFR and colorectal adenoma risk. Mutation research 782, 1–6 (2015).

38. Yang, X., et al. Gene body methylation can alter gene expression and is a therapeutic target in cancer. Cancer cell 26, 577–590 (2014).

39. Nakayama, T., et al. Epigenetic regulation of androgen receptor gene expression in human prostate cancers. Laboratory investigation; a journal of technical methods and pathology 80, 1789–1796 (2000).

40. Yang, M. & Park, J.Y. DNA methylation in promoter region as biomarkers in prostate cancer. Methods in molecular biology 863, 67–109 (2012).

41. Fan, M., et al. Diverse gene expression and DNA methylation profiles correlate with differential adaptation of breast cancer cells to the antiestrogens tamoxifen and fulvestrant. Cancer research 66, 11954–11966 (2006).

42. Heyn, H. & Esteller, M. DNA methylation profiling in the clinic: applications and challenges. Nature reviews. Genetics 13, 679–692 (2012).

43. Robertson, K.D. DNA methylation, methyltransferases, and cancer. Oncogene 20, 3139–3155 (2001).

44. Jung, M. & Pfeifer, G.P. Aging and DNA methylation. BMC biology 13, 7 (2015).

45. Cancer Genome Atlas, N. Comprehensive molecular portraits of human breast tumours. Nature 490, 61–70 (2012).

46. Cancer Genome Atlas Research, N. Comprehensive molecular characterization of clear cell renal cell carcinoma. Nature 499, 43–49 (2013).

47. Cancer Genome Atlas Research, N. Comprehensive molecular characterization of urothelial bladder carcinoma. Nature 507, 315–322 (2014).

48. Turcan, S., et al. IDH1 mutation is sufficient to establish the glioma hypermethylator phenotype. Nature 483, 479–483 (2012).

49. Jia, L., et al. Abnormally activated one-carbon metabolic pathway is associated with mtDNA hypermethylation and mitochondrial malfunction in the oocytes of polycystic gilt ovaries. Scientific reports 6, 19436 (2016).

50. Kaplan, E.L. & Meier, P. Nonparametric estimation from incomplete observations. Journal of American Statistical Association 53, 457–481 (1958).

51. Cerami, E., et al. The cBio cancer genomics portal: an open platform for exploring multidimensional cancer genomics data. Cancer discovery 2, 401–404 (2012).

52. Gentles, A.J., et al. The prognostic landscape of genes and infiltrating immune cells across human cancers. Nature medicine 21, 938–945 (2015).

53. Sproul, D., et al. Transcriptionally repressed genes become aberrantly methylated and distinguish tumors of different lineages in breast cancer. Proceedings of the National Academy of Sciences of the United States of America 108, 4364–4369(2011).

54. Carone, D.M. & Lawrence, J.B. Heterochromatin instability in cancer: from the Barr body to satellites and the nuclear periphery. Seminars in cancer biology 23, 99–108 (2013).

55. Katoh, Y., et al. Methionine adenosyltransferase II serves as a transcriptional corepressor of Maf oncoprotein. Molecular cell 41, 554–566 (2011).

56. Brosnan, J.T., da Silva, R.P. & Brosnan, M.E. The metabolic burden of creatine synthesis. Amino acids 40, 1325–1331 (2011).

## Online Methods References

24. Mehrmohamadi, M., Liu, X., Shestov, A.A. & Locasale, J.W. Characterization ofthe usage of the serine metabolic network in human cancer. Cell reports 9, 1507–1519 (2014).

29. Chen, X. & Ishwaran, H. Random forests for genomic data analysis. Genomics 99, 323–329 (2012).

32. Chen, E.Y., et al. Enrichr: interactive and collaborative HTML5 gene listenrichment analysis tool. BMC bioinformatics 14, 128 (2013).

42. Heyn, H. & Esteller, M. DNA methylation profiling in the clinic: applications andchallenges. Nature reviews. Genetics 13, 679–692 (2012).

51. Cerami, E., et al. The cBio cancer genomics portal: an open platform forexploring multidimensional cancer genomics data. Cancer discovery 2, 401–404(2012).

57. Zhu, Y., Qiu, P. & Ji, Y. TCGA-assembler: open-source software for retrieving and processing TCGA data. Nature methods 11, 599–600 (2014).

58. Ashburner, M., et al. Gene ontology: tool for the unification of biology. The Gene Ontology Consortium. Nature genetics 25, 25–29 (2000).

59. Marchal, C. & Miotto, B. Emerging concept in DNA methylation: role of transcription factors in shaping DNA methylation patterns. Journal of cellular physiology 230, 743–751 (2015).

60. Johnston, S.J. & Carroll, J.S. Transcription factors and chromatin proteins as therapeutic targets in cancer. Biochimica et biophysica acta 1855, 183–192 (2015).

61. Romero, P., et al. Computational prediction of human metabolic pathways from the complete human genome. Genome biology 6, R2 (2005).

62. Mermel, C.H., et al. GISTIC2.0 facilitates sensitive and confident localization of the targets of focal somatic copy-number alteration in human cancers. Genome biology 12, R41 (2011).

63. Costello, J.C., et al. A community effort to assess and improve drug sensitivity prediction algorithms. Nature biotechnology 32, 1202–1212 (2014).

64. Mentch, L. & Hooker, G. Quantifying Uncertainty in Random Forests via Confidence Intervals and Hypothesis Tests. The Journal of Machine Learning Research in press (2015).

65. Liaw, A. & Wiener, M. Classification and Regression by randomForest. R News 2, 18–22 (2002).

66. Friedman, J., Hastie, T. & Tibshirani, R. Regularization Paths for Generalized Linear Models via Coordinate Descent. Journal of statistical software 33, 1–22 (2010).

67. Lawrence, M., et al. Software for computing and annotating genomic ranges. PLoS computational biology 9, e1003118 (2013).

68. McLean, C.Y., et al. GREAT improves functional interpretation of cis-regulatory regions. Nature biotechnology 28, 495–501 (2010).

69. Brem, R.B., Storey, J.D., Whittle, J. & Kruglyak, L. Genetic interactions between polymorphisms that affect gene expression in yeast. Nature 436, 701–703 (2005).

70. Grosjean, P. & Ibanez, F. pastecs: Package for Analysis of Space-Time Ecological Series. R package version 1.3–18 (2014).

71. Tamborero, D., et al. Comprehensive identification of mutational cancer driver genes across 12 tumor types. Scientific reports 3, 2650 (2013).

72. Litovkin, K., et al. Methylation of PITX2, HOXD3, RASSF1 and TDRD1 predicts biochemical recurrence in high-risk prostate cancer. Journal of cancer research and clinical oncology 140, 1849–1861 (2014).

73. Gao, J., et al. Metscape: a Cytoscape plug-in for visualizing and interpreting metabolomic data in the context of human metabolic networks. Bioinformatics 26, 971–973 (2010).

74. Therneau, T. A Package for Survival Analysis in S_. version 2.38. (2015).

75. R-Core-Team. R: A Language and Environment for Statistical Computing.(2014).

76. Krzywinski, M., et al. Circos: an information aesthetic for comparative genomics. Genome research 19, 1639–1645 (2009).

